# The CoP Assay: A Robust Method for Quantitative Assessment of Food Intake in the Planarian *Dugesia japonica*

**DOI:** 10.64898/2026.07.22.740188

**Authors:** Reiichiro Fukushima, Yoshihiko Umesono

**Affiliations:** Graduate School of Science, University of Hyogo, 3-2-1 Kouto, Kamigori-cho, Ako-gun, Hyogo 678-1297, Japan

**Keywords:** Planarians, Food intake quantification, Food quality, Overconsumption, Satiety

## Abstract

Planarians exhibit a variety of behaviors in response to external stimuli and have emerged as a promising animal model for studying the neural regulation of behavior. Although their feeding behavior has been well documented, robust experimental methods for quantifying their food intake remain limited. In this study, we developed a novel methodology for the quantitative assessment of feeding termed the “Complete-or- Partial (CoP) assay.” This assay evaluates feeding responses as a binary outcome (complete consumption versus partial ingestion) in a volume-dependent manner at the individual level. Using this method, we examined the effects of food quality on feeding responses by comparing two distinct diets: a “basal diet” containing the minimum concentration of liver homogenate necessary to induce ingestion and support growth, and a “high-density diet” with a ten-fold higher concentration. The CoP assay enabled the robust estimation of food intake volume and successfully detected a significant overconsumption of the high-density diet compared to the basal diet, demonstrating that planarians regulate their food intake in a quality-dependent manner. Furthermore, individuals exhibiting partial ingestion did not immediately feed again even when presented with the same diet freshly prepared, indicating that they had reached satiety. Thus, we propose that the CoP assay offers a robust and scalable platform for evaluating both food intake volume and satiety status, providing a powerful tool to facilitate future molecular analyses of feeding regulation in planarians.

## INTRODUCTION

Feeding is a complex behavior essential for the survival of all animals and is regulated by a balance between two motivational states: hunger and satiety. Hunger signals promote food-seeking behavior and food intake, whereas satiety signals attenuate appetite to terminate feeding. An imbalance between these signals can lead to overconsumption, potentially resulting in metabolic disorders such as obesity, which has become a major public health concern in modern obesogenic environments (Mackenbach et al., 2014; Stuber et al., 2025). Therefore, a mechanistic understanding of the neural control systems governing the behavioral transition from hunger to satiety, particularly in response to food quality, is of paramount importance in basic neuroscience. To achieve this, the development of robust and quantitative methods is indispensable.

The freshwater planarian *Dugesia japonica* is a free-living invertebrate that possesses a simple central nervous system (CNS) (Agata et al., 1998; Tazaki et al., 1999; Umesono and Agata, 2009), but exhibits a variety of complex behaviors in response to external stimuli, including light, temperature, and food (Inoue et al., 2004, 2014, 2015; Shimoyama et al., 2016; Miyamoto et al., 2020). Consequently, it has emerged as a promising animal model for studying the neural regulation of behavior (Inoue and Agata, 2022). The feeding behavior of *D. japonica* is composed of a complex sequence of motor programs (Shimoyama et al., 2016; Miyamoto et al., 2020): i) chemotactic movements of individual animals toward a food source (e.g., liver homogenate), ii) protrusion of the pharynx out of the mouth in response to the food and subsequent extension of the pharynx toward it, iii) ingestion of the food into the gut through the distal opening of the pharynx, and iv) retraction of the pharynx into the body and movement away from the food source upon cessation of feeding. While feeding behavior is one of the best-studied behaviors in planarians, experimental methods for quantifying food intake remain limited. This is because measuring weight gain, the conventional approach to assessing food intake, is challenging due to the small size and aquatic nature of these organisms. Additionally, group-based measurement only provides the mean food intake of a population, which obscures individual variation and complicates statistical analysis. Although a quantitative analysis of food intake has been established in *D. japonica* (Mori et al., 2019; Miyamoto et al., 2020), further improvements in detection sensitivity are required for practical application. Therefore, establishment of a more sensitive method to overcome these limitations and quantify food intake at the individual level is highly desirable.

To address the limitations in quantifying planarian food intake, this study introduces the “Complete-or-Partial (CoP) assay,” a novel methodology that classifies individual feeding responses as a binary outcome (complete consumption or partial ingestion) in a volume-dependent manner. By applying this method to distinct basal and high-density dietary conditions, we demonstrated its utility for robustly estimating intake volume and detecting quality-dependent overconsumption. Furthermore, we validated that partial ingestion serves as a reliable indicator of satiety. Thus, we propose that the CoP assay, especially when combined with RNA interference, establishes a robust methodological platform for elucidating the neuronal mechanisms governing feeding regulation in planarians.

## MATERIALS AND METHODS

### Animals

A clonal strain of the planarian *D. japonica* (the HI strain) was used in this study. The animals were cultured at 22°C in autoclaved tap water. For all experiments, intact animals (7–8 mm in body length) were starved for at least 1 week in a culture container and then kept in a Petri dish for an additional 3 days prior to use.

### Diet preparation

A liver suspension was prepared by mixing beef liver homogenate with autoclaved tap water (1:2, v/v). A 2% agarose solution was prepared by dissolving 1 g of low-melting agarose powder in 50 mL of autoclaved tap water. An ink solution was prepared by diluting a PILOT Spotliter refill ink (SGRF-12SL-P; Pilot Corporation, Tokyo, Japan) at a 1:50 ratio with autoclaved tap water. Using these solutions, diet mixtures were prepared to a final volume of 40 µL, containing 5.0 μL of 2% agarose, 10 μL of ink solution, and the appropriate volume of liver suspension, with the remaining volume filled with autoclaved tap water. For the basal diet, 2.5 μL of liver suspension was used in the 40 μL of mixture, whereas for the high-density diet, 25 μL of liver suspension was added. These mixtures were aliquoted into appropriate volumes to form pellets on a piece of Parafilm using a micropipette and kept at <u>–</u>20℃ for 15<u>–</u>30 min to allow the agarose to gel.

### Cultivation with the basal diet

Each animal was placed into a well of a 12-well cell culture plate and supplied with 1.0 µL of a single basal diet pellet six times weekly. After each feeding, animals were observed under a stereomicroscope to confirm successful ingestion. Between feedings, the animals were transferred to and housed together in a 9-cm Petri dish.

### Measurement of body length

Photographs of individual animals with a straight posture were captured using a LEICA M165 FC microscope (Leica Microsystems, Wetzlar, Germany). For each imaging session, the animal was allowed to move freely in a 9-cm Petri dish. The body length of each animal was measured from at least five different photos using Fiji software, and the average value was defined as the body length of the individual.

### The CoP assay

Each animal was placed into a well of a 12-well cell culture plate. A single diet pellet of the appropriate volume was supplied to each well, and the animals were allowed to feed for 30 min. After the feeding period, the animals and pellets were observed under a stereomicroscope to assess feeding success. Feeding success was confirmed by the body expansion of individual animals, as well as the loss or deformation of the diet pellet. Animals with no remaining diet pellet in the well were classified as showing complete consumption (CC), whereas those with leftovers were classified as showing partial ingestion (PI). The rates of CC or PI for each group were calculated using the corresponding equation (Fig. 1C).

**Fig. 1.**
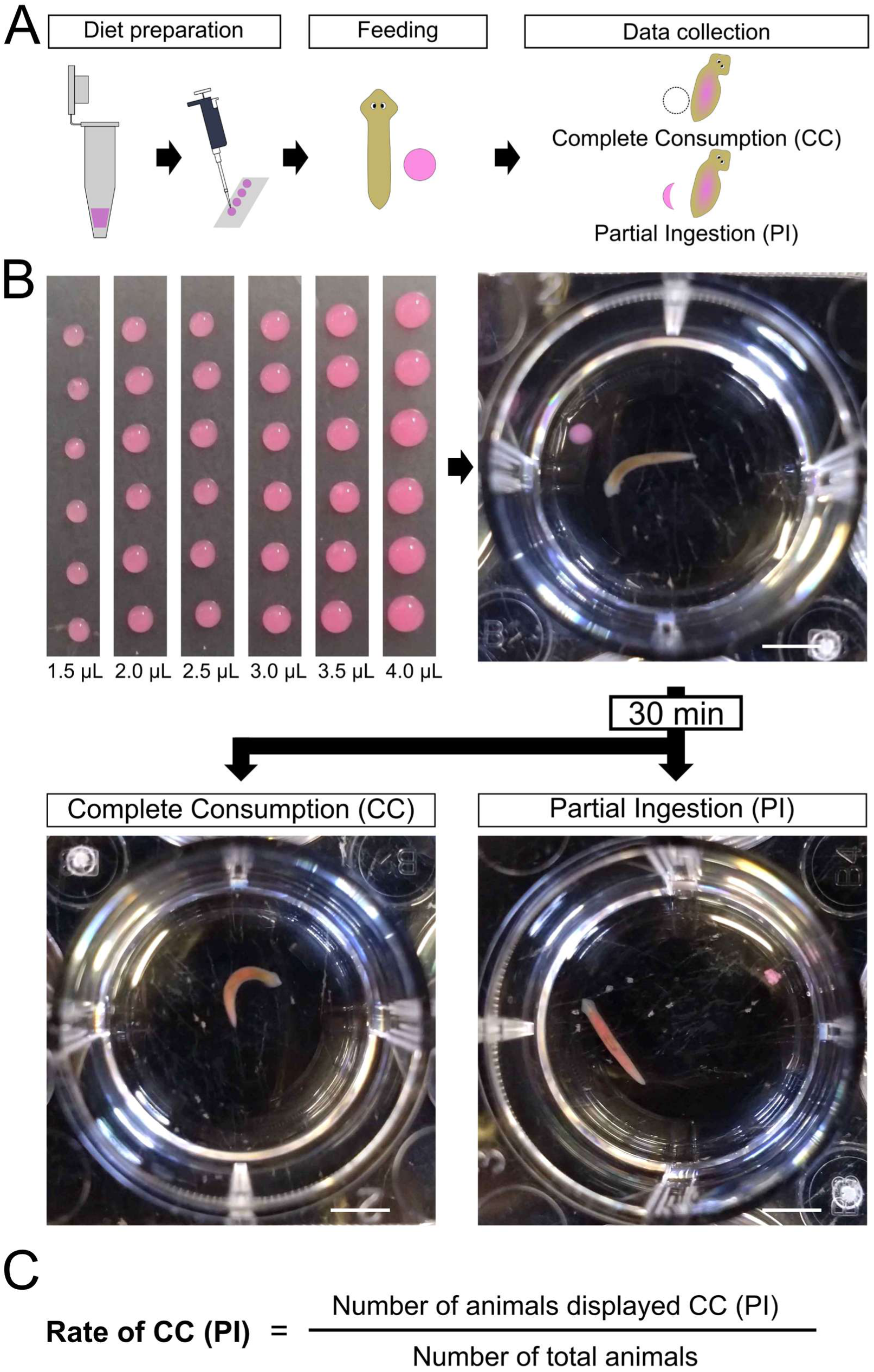
Methodology of the CoP assay. **(A)** Schematic drawing of the CoP assay. **(B)** Experimental procedure. Diet pellets with uniform volumes are prepared in a volume-dependent manner (upper left). Each animal is placed in a well of a 12-well plate and provided with a single diet pellet (upper right). After 30 min, an animal displaying complete consumption (CC) is shown (lower left), with no remaining diet in the well. Pink ink color is visible within the animal’s intestine as evidence of ingestion. An animal displaying partial ingestion (PI) is also shown (lower right), where ink color is observed in the intestine alongside the remaining diet pellet after the feeding period. **(C)** Formulae for CC and PI rates.

### Measurement of feeding duration

Animals were fed individually under the same conditions as in the CoP assay, and their behavior was recorded for 30 min against a white background using the camera application of a smartphone (Xperia 10 IV; Sony Corporation, Tokyo, Japan). The exact time points of feeding initiation and termination in each well were determined by reviewing the video recordings. Feeding initiation was defined as the moment when the animal positioned and anchored its tail tip near the diet. Feeding termination was defined as the detachment of the tail tip followed by the animal’s movement away from the diet. The feeding duration for each individual was calculated as the time interval between initiation and termination.

### Refeeding assay

Each animal was first fed with the initial diet pellet (prepared without the ink solution), and feeding success, as well as the occurrence of CC or PI, was assessed in the same manner as the CoP assay. Leftovers of the initial diet pellet were then removed, and the subsequent diet pellet containing the ink solution was immediately supplied without any interval. Subsequently, the success or failure of the second ingestion was evaluated under a stereomicroscope based on the presence of ink within the intestinal tract and the deformation of the second diet pellet.

### Statistical analysis

Data were analyzed using a one-way analysis of variance (ANOVA), a two-tailed Student’s *t*-test, or Fisher’s exact test, followed by a Bonferroni correction for multiple compariosns. A *p*-value of less than 0.05 (*p* < 0.05) was considered statistically significant in all tests.

## RESULTS

### The CoP assay: a novel methodology for the quantitative assessment of food intake

We developed a novel assay system termed the “Complete-or-Partial (CoP) assay,” in which each animal was supplied with a single diet pellet of varying volumes, and binary feeding outcomes, complete consumption (CC) or partial ingestion (PI), were recorded (for details, see Fig. 1 and MATERIALS AND METHODS). The proportion of CC relative to the total number of trials was calculated.

Next, we determined a “basal diet” that contains the minimum concentration of liver homogenate required to initiate ingestion and support growth. Using the composition used for feeding RNAi as a baseline (25 µL of liver suspension in a total volume of 40 µL; for details, see MATERIALS AND METHODS), we prepared diets in which the liver volume was gradually decreased in a stepwise manner. All animals tested in this study successfully ingested diets containing 2.5 µL (1/10 of the baseline) or more of liver suspension in a total volume of 40 µL. On the other hand, several animals failed to ingest diets containing 1.25 µL of liver suspension (Fig. 2A). Consequently, the composition of 2.5 µL of liver suspension in a total volume of 40 µL was defined as the basal diet.

**Fig. 2.**
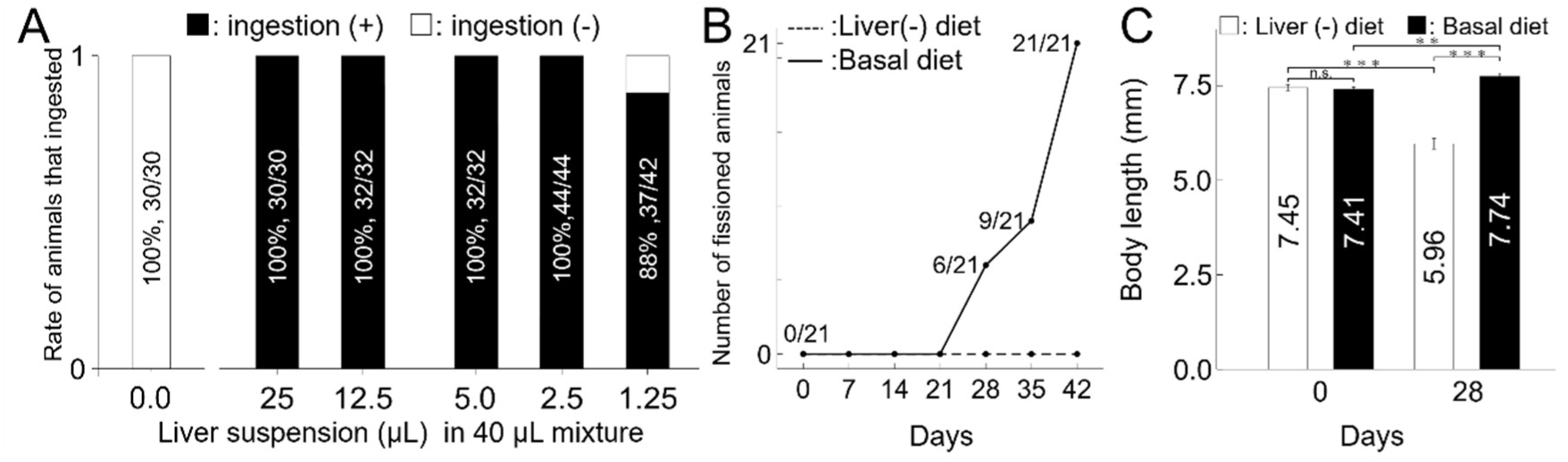
Establishment of the basal diet. **(A)** Feeding responses to diet pellets containing various volumes of liver suspension (ranging from 0.0 µL to 25 µL) in a total volume of 40 µL. **(B)** Number of animals that underwent fission during cultivation with the basal diet pellet or liver-free diet pellet. The solid line represents animals supplied with a single basal diet pellet prepared at a ratio of 2.5 µL liver suspension to 40 µL total mixture, and the dashed line represents those supplied with a liver-free diet pellet. Both diets were administered six times per week (*n* = 21 for each group). **(C)** Body length of animals on day 28 of cultivation. Data are shown for animals from **(B)** that did not undergo fission on the basal diet pellet by day 28 (*n* =15), and animals supplied with the liver-free pellet at day 28 (*n* = 21). Asterisks indicate statistical significance determined by a two-tailed Student’s *t*-test with Bonferroni correction (**: *P* < 0.01, \*\*\**P* < 0.001; n.s., not significant, *P* ≧ 0.05). Error bars represent means ± SEM.

We also confirmed that this basal diet is sufficient to support growth. Following 6 weeks of culture under this feeding condition (1.0 µL of a single basal diet pellet six times weekly), all animals underwent fission (21/21), serving as a definitive indicator of robust growth (Fig. 2B). Indeed, animals that did not undergo fission by day 28 of culture exhibited a significant increase in body length, whereas the control animals fed a liver-free diet (i.e., resulting in no ingestion) showed a significant decrease (Fig. 2C). Moreover, this disparity became even more pronounced when a larger amount of the basal diet was supplied in an independent experiment (Supplementary Fig. S1). These findings indicate that the basal diet provides adequate nutrition for both physiological maintenance and growth.

### The CoP assay reveals that food quality influences food intake

To examine whether food quality influences food intake using the CoP assay, we prepared a basal diet and a “high-density diet,” the latter containing a 10-fold higher concentration of liver suspension (i.e., identical in composition to the baseline). The volume of each diet supplied per individual animal ranged from 1.5 μL to 4.0 μL, at 0.5 µL increments. For both diets, the CC rate was 100% at 1.5 μL and 0% at 4.0 μL, indicating that the tested experimental range adequately covered the entire dynamic range (Fig. 3A, B). We found that the CoP assay enabled the robust estimation of food intake volume and successfully detected a significant overconsumption of the high-density diet compared to the basal diet (Fig. 3C). Furthermore, it was confirmed that the statistically significant difference between the two diets was robustly reproduced at specific pellet volumes (*n* = 3 independent experiments each for the 2.0 µL and 3.0 µL pellets; Supplementary Fig. S2).

**Fig. 3.**
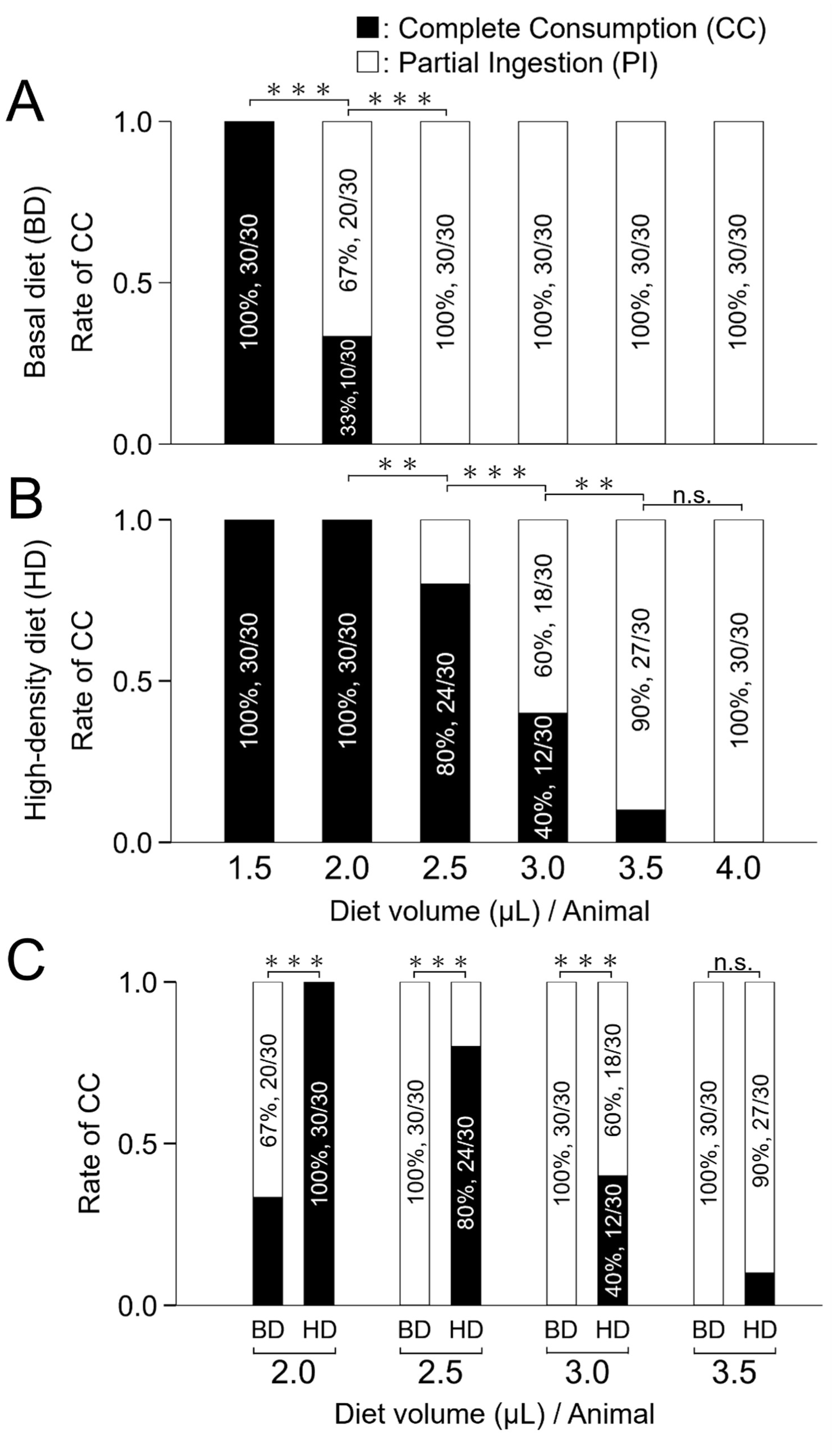
CoP assay. (A) Basal diet (BD). (B) High-density diet (HD). (C) Comparison of complete consumption rates between the BD and HD groups. For each experiment, 30 animals were used. Asterisks indicate statistical significance determined by Fisher’s exact test with Bonferroni correction (**: *P* < 0.01, ***: *P* < 0.001; n.s., not significant, *P* ≧ 0.05).

To investigate how the increased consumption is induced by the high-density diet, two types of measurements were conducted. First, to evaluate the difference in ingestion speed between the basal and the high-density diet, animals were individually supplied with 1.0 µL of either diet, and the duration of ingestion was measured. As expected, all tested animals displayed CC. We found that the time required to completely consume the high-density diet was significantly shorter than that for the basal diet (Fig. 4A). Second, to evaluate the difference in the time required to terminate feeding between the two diets, animals were individually supplied with 5.5 µL of either diet, and their feeding duration was measured. As expected, all tested animals displayed PI. We found that the duration of ingestion for the high-density diet was significantly longer than that for the basal diet (Fig. 4B).

**Fig. 4.**
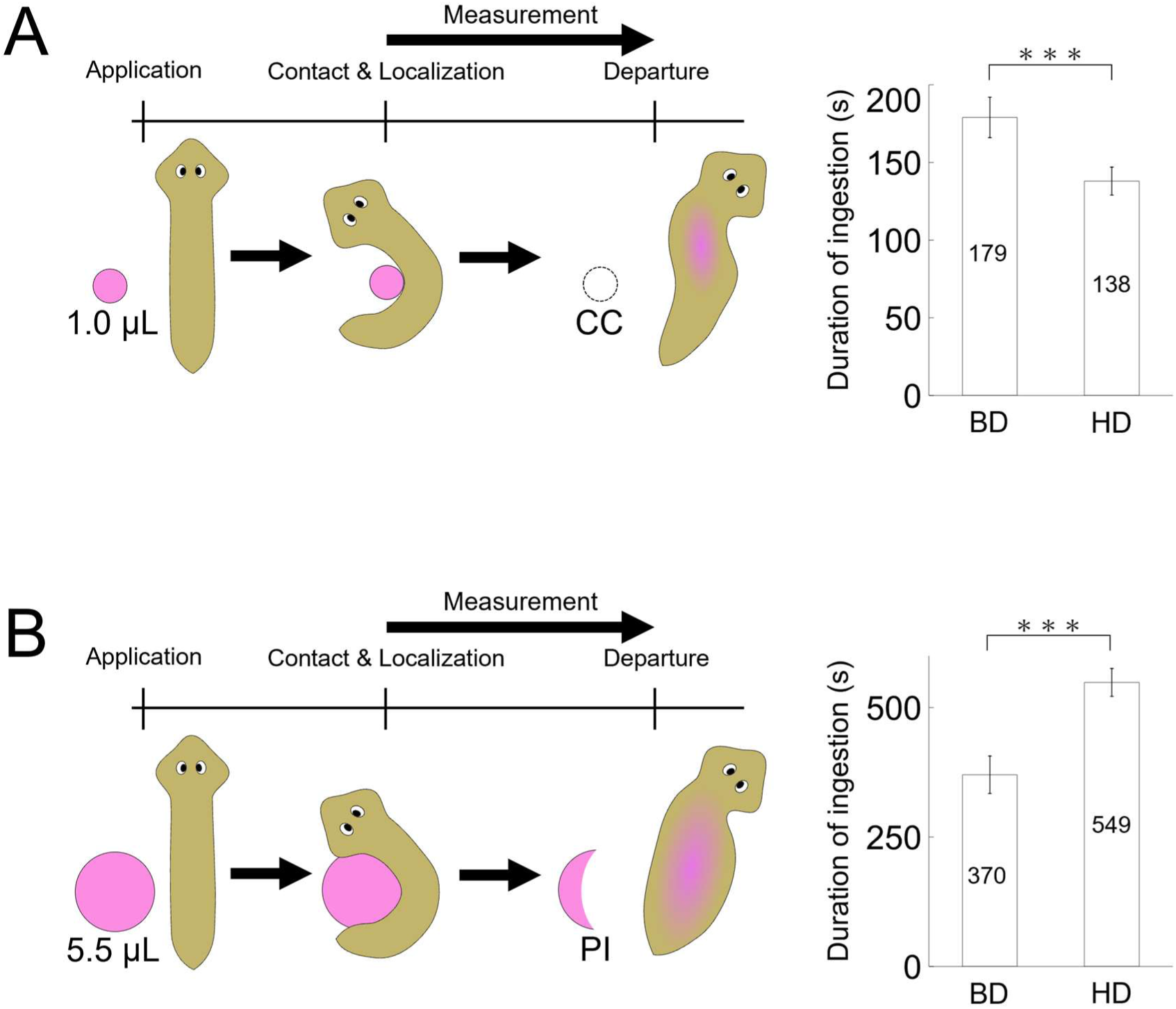
Comparison of feeding durations between the basal and high-density diets. **(A)** Duration for complete consumption (CC) of 1.0 µL of a single diet pellet. (Left) Schematic drawing of the experimental setup. (Right) Mean consumption times (*n* = 7 for the basal diet; *n* = 12 for the high-density diet). **(B)** Duration for partial consumption (PI) of 5.5 µL of a single diet pellet. (Left) Schematic drawing of the experimental setup. (Right) Mean consumption times (*n* = 10 for the basal diet; *n* = 12 for the high-density diet). Error bars represent mean ± SEM. Asterisks indicate statistical significance determined by Student’s *t*-test (***: *P* < 0.001).

These findings demonstrate that the overconsumption of the high-density diet is driven by both the acceleration of feeding speed and the extension of ingestion duration.

### The CoP assay effectively captures the onset of satiety

We developed a “refeeding assay” to investigate the relationship between PI and the onset of satiety. Individual animals were supplied with 5.5 µL of diet in two consecutive 30-minute trials without an inter-trial interval (Fig. 5A). In the primary trial, all tested animals (30/30) displayed PI, which was consistent with our previous findings (Fig. 4) and validated the experimental conditions. In the secondary trial, however, no animals showed any further ingestion (30/30), regardless of whether they were presented with a fresh basal or high-density diet (Fig. 5B, C). These findings demonstrate that PI in the CoP assay reliably reflects the onset of satiety in planarians.

**Fig. 5.**
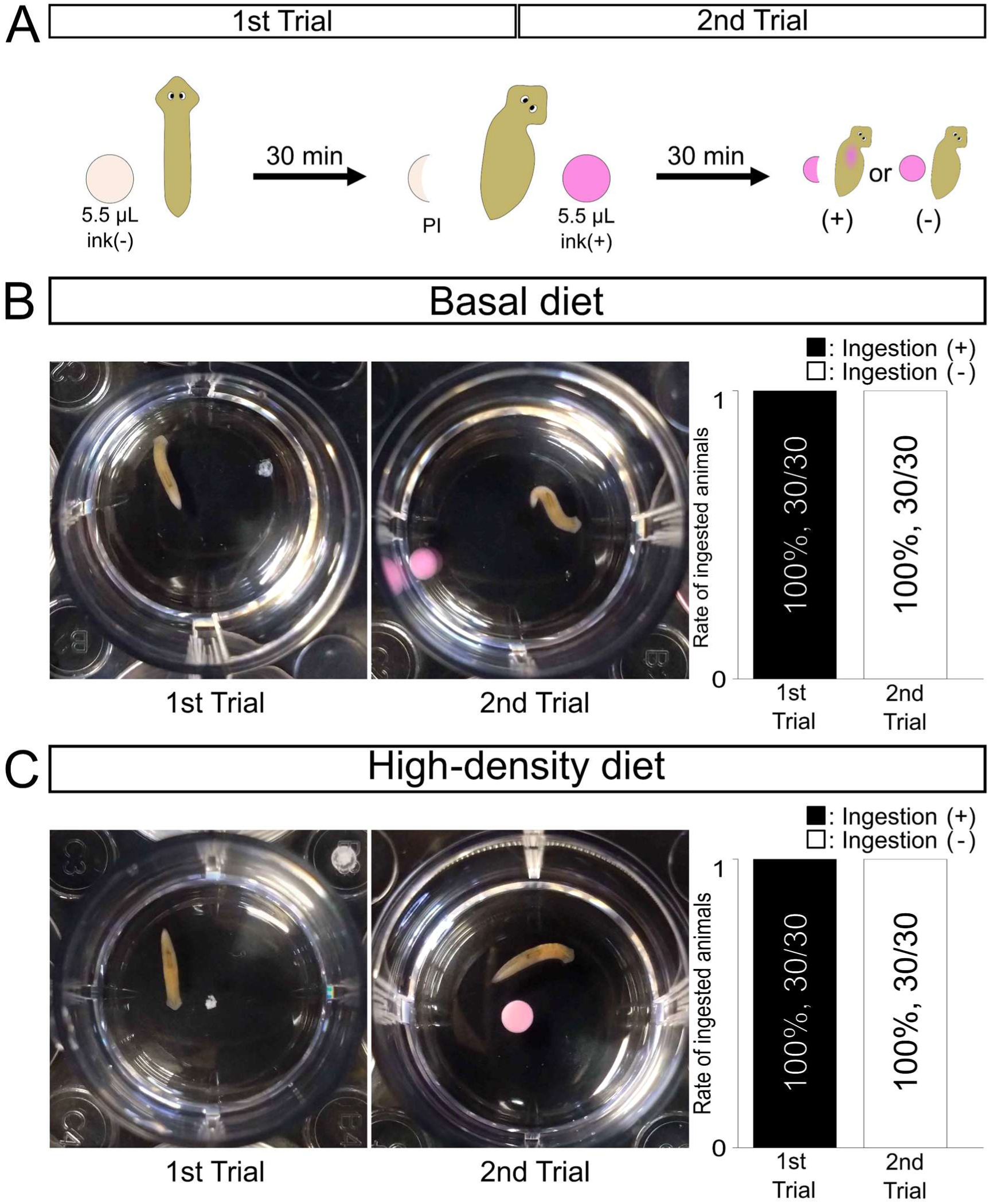
Refeeding assay on the basal diet and high-density diet. **(A)** Schematic drawing of the refeeding assay. **(B)** Refeeding assay on the basal diet. (Left panel) An animal supplied with a single basal diet pellet (5.5 µL) in a well of a 12-well plate displaying partial ingestion (PI) after 30 min, with the remaining ink-free diet observed after ingestion. (Center panel) Immediately afterward, without an interval, the animal is supplied with another fresh single basal diet pellet (5.5 µL) containing pink ink color. After an additional 30 min, the animal displaying no feeding is shown, where the second pellet remains completely intact and no ink color is visible within the animal’s intestine. (Right panel) The graph displays the success rate of ingestion, where black and white bars represent animals that succeeded and failed to ingest, respectively. **(C)** Refeeding assay on the high-density diet. For each experiment, 30 animals were used.

Taken together, our findings demonstrate that planarians precisely regulate their food intake in a food quality-dependent manner, a process tightly linked with the transition from hunger to satiety.

## DISCUSSION

The CoP assay offers a practical platform to indirectly yet robustly estimate the maximum food intake capacity at the individual level by strictly controlling the food volume provided. Thus, this methodology effectively overcomes the limitations of conventional population-based measurements. One of the most significant advantages of this assay is its quantitative resolution, which enables the detection of dynamic behavioral transitions, such as the shift from complete consumption to partial ingestion, indicating the onset of satiety. In particular, the “basal diet” developed in this study serves as a standardized physiological baseline for individual animals (Fig. 2). This condition is ideal for future screening studies aimed at identifying chemical substances that promote or inhibit feeding behavior. For example, by applying the CoP assay to the basal diet supplemented with various sensory cues (such as odors and tastants) or metabolic modulators, we can systematically screen for and identify the bioactive molecules underlying the enhanced feeding motivation that drives overconsumption, which may be enriched in high-density diets. Furthermore, when combined with RNA interference targeting specific candidate genes (Takano et al., 2007; Rouhana et al., 2013, Hattori et al., 2018), this assay will enable us to pinpoint the neuronal circuits that integrate food quality and metabolic states to regulate feeding responses. Indeed, we are currently utilizing the CoP assay to analyze the functions of specific neuronal cell types during feeding, and have already confirmed the robustness of this assay for statistical comparison between distinct experimental groups with an optimized diet pellet volume (data not shown). These ongoing efforts underscore the practical advantage of the CoP assay for dissecting the neural circuits governing planarian feeding behavior.

Finally, the core principle of the CoP assay is highly adaptable to a wide range of small aquatic invertebrates. By scaling the experimental parameters of food volume to suit the target species, this methodology can serve as a versatile platform to examine feeding events across aquatic ecology, ecotoxicology, and comparative physiology.

## Supporting information

Fukushima_Supplementary_Figures

## ACKNOWLEDGMENTS

During the preparation of this manuscript, the authors used Gemini (Google) to improve the English language and phrasing. After using this service, the authors reviewed and edited the content as needed and take full responsibility for the final version of the manuscript. We thank Dr. Elizabeth Nakajima for editing and proofreading the manuscript.

## AUTHOR CONTRIBUTIONS

YU and RF designed the study. RF performed all experiments. YU and RF interpreted the results and prepared the manuscript. RF also prepared all the figures. Both authors read and approved the final manuscript.

---

**Supplementary Fig. S1.** Comparison of fission rate and body length between the basal and liver-free diet groups. **(A)** Number of animals that underwent fission during cultivation with the basal diet pellet or liver-free diet pellet. The solid line represents animals supplied with an excess amount of the basal diet pellet prepared at a ratio of 2.5 µL liver suspension to 40 µL total mixture, and the dashed line represents those supplied with the same amount of the liver-free diet pellet. Both diets were administered six times per week (*n* = 21 for each group). **(B)** Body length of animals on day 28 of cultivation. Data are shown for animals from **(A)** that did not undergo fission on the basal diet pellet by day 28 (*n* = 16), and animals supplied with the liver-free diet pellet at day 28 (*n* = 21). Asterisks indicate statistical significance determined by a two-tailed Student’s *t*-test with Bonferroni correction (\*\*\**P* < 0.001; n.s., not significant, *P* ≧ 0.05). Error bars represent means ± SEM.

**Supplementary Fig. S2.** Comparison of complete consumption between the basal diet (BD) and high-density diet (HD) groups determined by the CoP assay. Two independent assays were performed and provided reproducible results. For each experiment, 30 animals were used. Asterisks indicate statistical significance determined by Fisher’s exact test with Bonferroni correction (**: *P* < 0.01, ***: *P* < 0.001).

