## Supplementary material for "The CoP Assay: A Robust Method for Quantitative Assessment of Food Intake in the Planarian *Dugesia japonica*": Fukushima_Supplementary_Figures

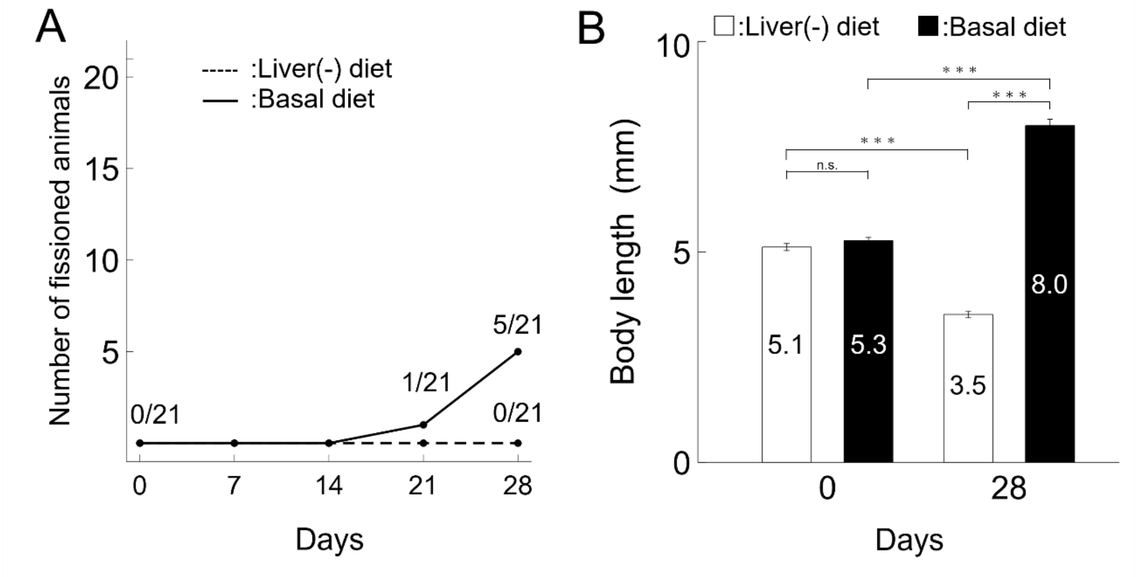


**Supplementary Fig. S1.** Comparison of fission rate and body length between the basal and liver-free diet groups. **(A)** Number of animals that underwent fission during cultivation with the basal diet pellet or liver-free diet pellet. The solid line represents animals supplied with an excess amount of the basal diet pellet prepared at a ratio of 2.5 μL liver suspension to 40 μL total mixture, and the dashed line represents those supplied with the same amount of the liver-free diet pellet. Both diets were administered six times per week ($n = 21$ for each group). **(B)** Body length of animals on day 28 of cultivation. Data are shown for animals from **(A)** that did not undergo fission on the basal diet pellet by day 28 ($n = 16$), and animals supplied with the liver-free diet pellet at day 28 ($n = 21$). Asterisks indicate statistical significance determined by a two-tailed Student's *t*-test with Bonferroni correction (***$P < 0.001$; n.s., not significant, $P \geqq 0.05$). Error bars represent means ± SEM.

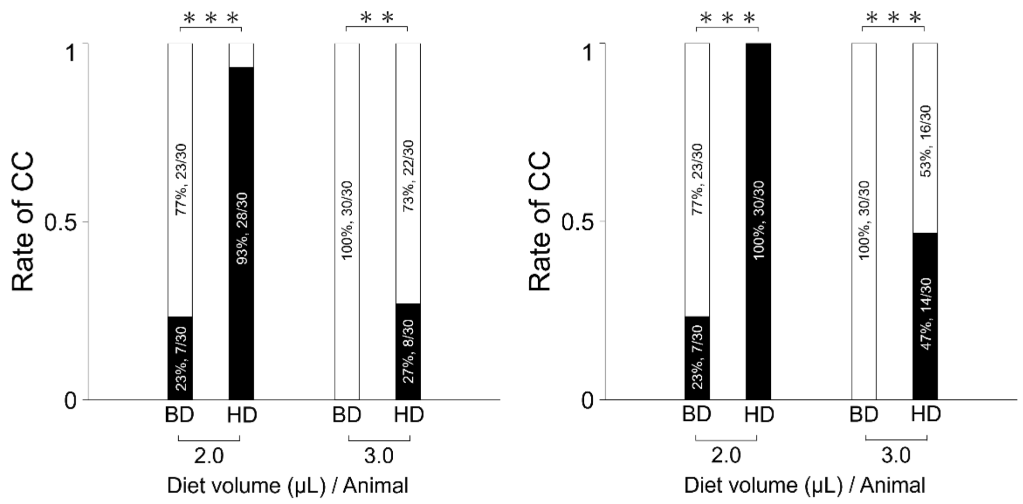


**Supplementary Fig. S2.** Comparison of complete consumption between the basal diet (BD) and high-density diet (HD) groups determined by the CoP assay. Two independent assays were performed and provided reproducible results. For each experiment, 30 animals were used. Asterisks indicate statistical significance determined by Fisher's exact test with Bonferroni correction (**: $P < 0.01$, ***: $P < 0.001$).
